# Transcriptomic and proteomic dynamics during metamorphosis of Pacific oyster *Crassostrea gigas*

**DOI:** 10.1101/2020.03.25.004614

**Authors:** Fei Xu, Guofan Zhang

## Abstract

Many marine invertebrate phyla are characterized by indirect development. These animals transit from planktonic larvae to benthic adults via settlement and metamorphosis, which contributes to the adaption to the marine environment. Studying the biological process of metamorphosis is thus a key to understand the origin and evolution of indirect development. Numerous studies have been conducted on the relationships of metamorphosis with the marine environment, microorganisms, as well as the neurohormones, however, little is known on the gene regulation network (GRN) dynamics during metamorphosis. Metamorphosis competent pediveliger of Pacific oyster *Crassostrea gigas* was assayed in this study. By identifying genes enriched in competent pediveliger and early spat, as well as pediveligers treated with epinephrine, the dynamics of genes and proteins was examined with transcriptomics and proteomics methods. The results indicated significantly different gene regulation networks before, during, and post metamorphosis. Genes encoding membrane integrated receptors and related to the remodeling of the nervous system were upregulated before the initiation of metamorphosis. Massive biogenesis, e.g., various enzymes and structural proteins, occurred during metamorphosis. Correspondingly, the protein synthesis system was comprehensively activated after epinephrine stimulation. Hierarchical downstream gene networks were also stimulated, where some transcription factors showed different temporal response patterns, including some important Homeobox, basic helix-loop-helix factors and nuclear receptors. Nuclear receptors, as well as their retinoic acid receptor partners, should play critical roles during the oyster metamorphosis, although they may not be responsible for the initiation process. Enriched genes in early spat were mainly related to environmental stress responses, indicating the GRN complexity of the transition stage during oyster metamorphosis.

## Introduction

Knowledge of vertebrates and arthropods model animals, which are primitively direct developers (Davidson et al., 1995, Sly et al., 2003), are far from enough to understand the evolution of metazoans development fully (Xu et al., 2016). Studies on the gene regulation network of transition stage of “maximal” indirect developers (Davidson et al., 1995), a large group of marine organisms forming larvae during development, will provide insight into the evolutionary mechanism of animals ontogeny (Raff, 2008). Indirect developers commonly have a transition stage from planktonic larvae to benthic adults, where some fundamental bioprocesses occur, such as settlement and metamorphosis. Furthermore, the transition usually undergoes rapidly (Hadfield, 2000). As the adult body plan is distinct from larvae, understanding of how the gene regulation network (GRN) is re-organized is the key to gain insight into the complexity of the transition (Heyland et al., 2006).

While the metamorphosis GRN mechanism of model animals, e.g., frog and fruit fly, has been widely studied, marine indirect developers did not receive sufficient attention. A single hormone usually plays an important role in the initiation of the metamorphosis of frog (thyroid hormone, TH) (Brown et al., 2007) and fruitfly (ecdysone), which induces diverse phenotypic changes and intracellular responses. Thyroid hormone induces GRN in tadpoles through activating transcription factor thyroid receptors (TRs), together with their retinoic acid receptor (RXR) partners (Shi et al., 2012). Injection of ecdysone induces the puffing of salivary gland chromosomes, resulting in transcription activation of many genes, and initiation of *Drosophila* molting (Hill et al., 2013). Ecdysone receptor (EcR), as well as its partner USP (ultraspiracle, an ortholog of vertebrate RXR), plays a central role during this biology progress.

Comparing to the profound studies of model animals from Deuterostomia and Ecdysozoa, the genetic background of the third main branch of Bilateria, Lophotrochozoans, is poorly understood. Recent studies suggested that Lophotrochozoan species may contain integrated hormone systems. Important physiological roles of some various hormones, such as ecdysone (Lafont et al., 2009), TH (Huang et al., 2015), octopamine (Ji et al., 2016), and epinephrine (Coon et al., 1986) were all reported in Lophotrochozoans. Furthermore, Lophotrochozoans may maintain the most comprehensive hormone regulatory system compared with vertebrates and insects. For example, molluscs use both epinephrine and octopamine, while vertebrates make extensive use of epinephrine but few of octopamine, and insects use OA but not epinephrine (Adamo, 2008, Ji et al., 2016). Genes related to both thyroid hormone and ecdysone system (synthesis enzymes, TR, EcR, and RXR) have been identified in Pacific oyster *Crassostrea gigas* (Zhang et al., 2012, Huang et al., 2015). Indeed, both vertebrates and insects are thought to be highly specialized phyla at the view of developmental regulatory mechanisms (Peterson et al., 2000, Luo et al., 2018, Guijarro-Clarke et al., 2020). In this view, Lophotrochozoans represent the most general mode of Bilaterian development (Davidson et al., 1995). Lophotrochozoan development GRNs characterization is in badly need, although representatives from the other two Bilateria branches have been studied comprehensively.

Pacific oyster *Crassostrea gigas* is a typical indirect developer, and one of the well-studied Lophotrochozoans because of its biology and aquaculture importance (Hedgecock et al., 2005). The planktonic larvae stage usually last two weeks and then form metamorphosis competent pediveliger at a size larger than 300 μm. Pediveligers detect substratum with their foot. Settlement and metamorphosis are then induced by the proper environment, chemicals, or biofilm. Some neurotransmitters, such as epinephrine, can induce direct metamorphosis of pediveliger and have thus been successfully used to produce cultchless oyster in hatcheries (Coon et al., 1986). As oyster settlement and metamorphosis are quick and usually nonsynchronous, it is hard to collect transitioning samples in real-time. Epinephrine induced pediveligers provided an excellent model to study the GRN during the metamorphosis stage. We collected both epinephrine induced samples and naturally attached (indicating successful settlement and metamorphosis) samples to conduct transcriptome and proteomics studies. In this way, the GRN complexity of the transition stage was revealed by comparing genes expression patterns before and post metamorphosis.

## Materials and methods

### Oyster treatment and sampling

Larvae production and culture of Pacific oyster *Crassostrea gigas* was conducted at the Laodong Aquaculture Breeding Company, Qingdao, China, as described previously (Zhang et al., 2012). In brief, gametes of gonad matured oysters were dissected and washed before insemination to produce massively mated offspring. Zygotes incubation and following larvae culture were conducted in 25 m^3^ of sand-filtered seawater at 25°C, with salinity around 30. After 20 days of culture, metamorphosis competent larvae (pediveliger) were harvested and maintained in 5 L seawater at the concentration of 10 per mL for epinephrine treatment. Epinephrine hydrochloride (EPI, Solarbio, China) solution was added at the final concentration of 1×10^−4^ M (EPI experiment). After 20 min of treatment, pediveligers were washed and transferred into fresh seawater. General sampling time points were set as: 20 min post-treatment (i.e., sampling immediately after wash), 3 h, 6 h, 12 h and 24 h. At each sampling time point, pediveligers were collected and immediately frozen in liquid nitrogen before being stored at -80°C. Pediveligers without any treatment were sampled in parallel as controls. The exact sampling time for three experiments (Exp) were: 20 min post-treatment (mpt) and 6 h post-treatment (hpt) for Exp#1, 20 mpt, 3 hpt, 6 hpt, 12 hpt for Exp#2, 20 mpt, 3 hpt, 6 hpt, 12 hpt, 24 hpt for Exp#3.

In another experiment (Exp#4, spat experiment), metamorphosis competent pediveligers were sampled in successive five days; attached spats were then sampled in the next consecutive five days. In this way, five pediveliger samples and five early spat samples were collected to compare the gene expression differences before and after metamorphosis.

### RNA extraction, sequencing, and data analysis

RNA was extracted using TRIzol according to the manufacturer’s protocol. Single-end 49 bp (Exp #1), 50 bp (Exp#2 & #3), or 70 bp (Exp#4) RNA sequencing (RNAseq) was performed on Illumina HiSeq 2000 platform (Illumina, Inc., San Diego, CA, USA) from libraries construed as has been described in previous report (Zhang et al., 2012). Cleaned sequencing reads were mapped to the oyster genome with TopHat (Trapnell et al., 2009). Gene expression levels were quantified as fragments per kilobase per million mapped reads (FPKM) with the Cufflinks (Mortazavi et al., 2008). Functional annotations of the proteins were conducted using the Blast2GO program against the NCBI non-redundant protein database (Gotz et al., 2008).

Genes specifically expressed during pediveliger stage were identified by analysing the published developmental transcriptome data (Zhang et al., 2012). Pediveliger specifically enriched gene was defined as the max FPKM value was at pediveliger stages, while the values of samples before the umbo larvae stage were less than 5, and the values of the juvenile were less than the half of the max value. At the same time, FPKM values from the late umbo stage to the pediveliger should fit a line curve with a positive correlation (R^2^>0.8 with a positive slope value).

R software (Ihaka et al., 1996) was used for statistics and plotting, in which, rgl package was used to draw the principal component analysis (PCA) result, DEGseq package (Wang et al., 2010) was used to identify differentially expressed genes (DEGs), clusterProfiler package (Yu et al., 2012) was used for KEGG enrichment. Blast2go was used for gene ontology (GO) enrichment, where only the most specific significant terms were kept. During DEGs analysis, data from 20 mpt samples were not used, as PCA analysis indicated that gene expression patterns for 20 mpt samples did not differentiate from control samples. In this way, the epinephrine treated sample and its control at 6 hpt in Exp#1 were compared directly with the non-replication strategy. Epinephrine treated samples and their controls at 3 hpt, 6 hpt, 12 hpt in Exp#2 were regarded as two groups of samples with three biology replications, samples at 3 hpt, 6 hpt, 12 hpt, and 24 hpt in Exp#3 were regarded as two groups with four biology replications. Up-regulated genes (URGs) and down-regulated genes (DRGs) were then integrated by intersecting the DEGs from the three experiments. As the power of non-replication strategy is limited in Exp#1, we also consider the intersected DEGs from Exp#2 and Exp#3 in some analysis if the general expression pattern (up/down compared with control) did not conflict with the result from Exp#1. In the spat experiment, pediveliger samples and spat samples were treated as two groups (each with five biological replications) to determine the significance (q-value) of gene expression difference. DEGs were further defined as those having two-fold differences for both the maximum and average FPKM values between the two group samples.

R package WGCNA (Langfelder et al., 2008) was used to conduct weighted correlation analysis. Boolean values were assigned as sample traits in the EPI experiment (0 for control samples and 20 mpt samples, 1 for other EPI treated samples) and spat experiment (0 for pediveliger samples and 1 for spat samples). Correlations were then calculated between gene expression pattern and sample traits. Multiple testing P values were adjusted according to Benjamini & Hochberg method (Benjamini et al., 1995).

### Protein extraction, sequencing, and analysis

Methods used for protein experiments were as have been described in the previous report (Qiao et al., 2012, Meng et al., 2017) and were slightly modified. In brief, proteins were extracted from the pediveligers of 3 h, 12 h and 12 h control groups in Exp#2 & #3 with lysis buffer for following treatment. The eluted peptides were monitored at 214 nm wavelength and pooled into 12 fractions. Each fraction was vacuum-dried after being desalted with the Strata-X C18 column (Phenomenex). Each fraction was resuspended and separated on an LC-20AD Nano HPLC (Shimadzu, Kyoto, Japan) by the autosampler onto a 2 cm C18 trap column. Data acquisition was performed using a TripleTOF^®^ 6600 system (Sciex, Concord, ON, Canada). The MS data were converted into MGF files using Proteome Discoverer™ 1.2 (Thermo Fisher). Protein identification and quantification were performed with Mascot 2.3.02 software (Matrix Science, London, United Kingdom). Genome sequences and annotations were downloaded from NCBI. Functional annotations of the proteins were conducted using the Blast2GO program against the NCBI non-redundant protein database.

## Results

### Specifically enriched genes during the pediveliger stage

A total of 389 genes were identified to be specifically enriched during the pediveliger stage. Cellular component terms “integral component of membrane (GO:0016021)”, “extracellular region (GO:0005576)”, “plasma membrane (GO:0005886)”, and “postsynapse (GO:0098794)” were significantly enriched. Related biological process terms included: “G protein-coupled receptor signaling pathway (GO:0007186)”, and “excitatory postsynaptic potential (GO:0060079)” (Supplementary FigureS1). Two main groups of receptors were enriched in this gene set: G protein-coupled receptors (GPCRs) and nicotinic acetylcholine receptors (nAChRs). Pediveliger specifically expressed G protein-coupled receptors (GPCRs) included putative *octopamine receptor* (CGI_10017568), *gamma-aminobutyric acid receptor* (CGI_10018562), *dopamine receptor* (CGI_10012064), and some neuropeptide receptors (CGI_10027017, CGI_10006524, CGI_10007542, CGI_10013409). Other types of receptors included possible nicotinic acetylcholine receptors (nAChRs, CGI_10027928, CGI_10011051, CGI_10017000, CGI_10009630, CGI_10015878, CGI_10008996, CGI_10015877), glutamate receptors (CGI_10021967, CGI_10028046), *glycine receptor* (CGI_10001715), toll-like receptors (CGI_10020169, CGI_10021737, CGI_10021741), and *purinergic receptor* (CGI_10019725). Many other genes not well annotated also showed specifically high express level during pediveliger, e.g., CGI_10003237, CGI_10004853, CGI_10005634, CGI_10026725, CGI_10007887, CGI_10004748, CGI_10009961, etc. Some genes related to the formation of the adult nervous system were also upregulated during the pediveliger stage, such as SCO-spondins (CGI_10022619 & CGI_10022620), neuroligins (CGI_10007153, CGI_10007154), *neurotrimin* (CGI_10014425) and *tenascin* (CGI_10002258).

### Enriched genes in early spat post attachment

A total of 1390 upregulated genes (URGs) and 373 down-regulated genes (DRGs) were identified in the attached spat. GO enrichment analysis indicated that genes related to biological processes “oxidation-reduction process (GO:0055114)”, “negative regulation of endopeptidase activity (GO:0010951)”, “chitin catabolic process (GO:0006032)”, and “removal of superoxide radicals (GO:0019430)” was significantly enriched in the URGs. Another group of genes enriched in the URGs was structural protein genes (Collagens and Chitins). GO enrichment analysis identified cellular component and molecular function terms “collagen trimer (GO:0005581)”, “extracellular matrix (GO:0031012)” and “chitin binding (GO:0008061)”. Correspondingly, KEGG enrichment showed that genes related to pathways “neuroactive ligand-receptor interaction (map04080)” (Supplementary FigureS2) and “ECM-receptor interaction (map04512)” were upregulated. No enriched GO term was identified in DRGs.

### Morphologic and transcriptomic response of pediveligers to epinephrine

Pediveligers showed quick morphologic response to epinephrine. Larva transformed from planktonic to benthic immediately after the treatment. The calcified shell was observed from samples collected since 6 hpt (Figure 1a). Principal component analysis (PCA) was conducted to determine the gene expression pattern differences between control and EPI treatment groups (EPI experiment). The result indicated that EPI treatment groups and control groups were distinctly separated, while three biological replicates were also distinguishable, suggesting significant genomic response of pediveliger to epinephrine stimulation (Figure 1b). However, treatment groups 20 mpt were clustered together with control groups.

**Figure 1.**
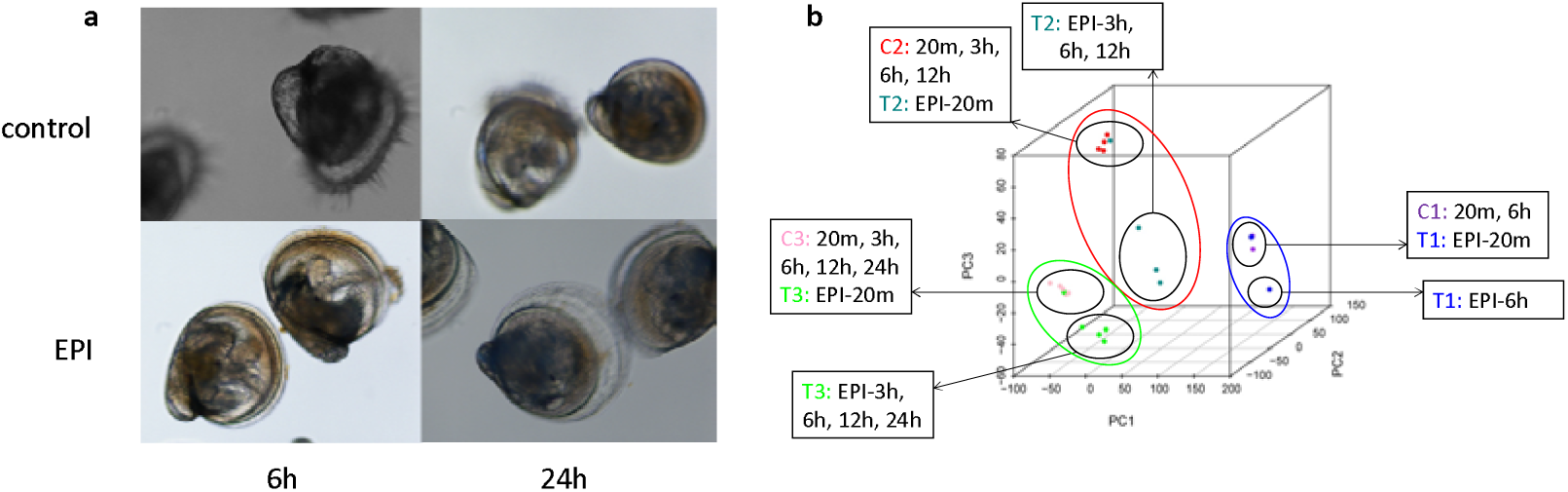
The morphologic change of Pacific oyster competent pediveligers after epinephrine stimulation (a) and the principal component analysis on the RNAseq data (b). Calcified shell grew quickly post epinephrine stimulation. The first RNAseq replication is shown with purple (C1:20m, 6 h, controls at 20 min and 6 h) and blue (T1: EPI-20m, EPI-6h, EPI treated groups at 20 min and 6 h) dots. The second replication is shown with red (C2:20m, 3h, 6h, 12h, controls at 20 min, 3 h, 6 h and 12 h) and dark green (T2: EPI-20m, EPI-3h, EPI-6h, EPI-12h, EPI treated groups at 20 min, 3 h, 6 h and 12 h) dots. The third replication is shown with pink (C3:20m, 3h, 6h, 12h, 24h, controls at 20 min, 3 h, 6 h, 12 h and 24 h) and blue (T2: EPI-20m, EPI-3h, EPI-6h, EPI-12h, EPI-24h, EPI treated groups at 20 min, 3 h, 6 h, 12 h and 24 h) dots. The EPI treated groups are separated from the controls in all the three replications, except for the 20 min groups clustered with the controls.

A total of 1633 DEGs (including 825 URGs and 808 DRGs) were identified by comparing metamorphosis competent pediveligers and those treated with epinephrine. GO enrichment analysis on URGs indicated that terms were mainly related with gene transcription, protein biogenesis and transport such as “regulation of transcription, DNA-templated (GO:0006355)”, “regulation of translational initiation (GO:0006446)”, “protein folding (GO:0006457)”, and “COPII-coated vesicle budding (GO:0090114)” (Supplementary Figure S3). These were further supported by KEGG enrichment analysis, where pathways related with “Protein processing in endoplasmic reticulum (map04141)”, “Ribosome biogenesis in eukaryotes (map03008)”, “RNA transport (map03013)”, and “Aminoacyl-tRNA biosynthesis (map00970)” were enriched in URGs. Main related genes included five translation initiation factors (*eIF1*, CGI_10007072; *eIF1A*, CGI_10026140; *eIF2*, CGI_10028301, CGI_10003139, CGI_10010423; *eIF3*, CGI_10010150, CGI_10024813, CGI_10008836, CGI_10005256, CGI_10006002, CGI_10027107; *eIF4E*, CGI_10000487; *eIF4G*, CGI_10022856 and *eIF5*, CGI_10013849), six molecular chaperones (CGI_10017621, CGI_10016162, CGI_10011081, CGI_10025730, CGI_10028542, CGI_10009495), and all subunits composing chaperonin proteins group I (*GroEL*, CGI_10011081 & *GroES*, CGI_10011080) and group II (subunit alpha CGI_10018918; beta, CGI_10026913; gamma, CGI_10004346; delta, CGI_10017263; epsilon, CGI_10015118; zeta, CGI_10004395; eta, CGI_10013567; theta, CGI_10002375) (Figure 2a&b). Genes related with intracellular receptor signalling pathway were also enriched: “steroid hormone mediated signaling pathway (GO:0043401)” including nuclear receptors (CGI_10003107, CGI_10007403, CGI_10011713, CGI_10020896), together with their possible interacting partners *retinoid X receptor* (RXR, CGI_10004075) and *retinoic acid receptor* (RAR, CGI_10028545). Expression level of several homeobox and basic helix-loop-helix (bHLH) transcription factors were upregulated, including *Cers* (CGI_10021077), *Hox1* (CGI_10024083), *Meis* (CGI_10019589), *Mkx* (CGI_10011802), *Tgif* (CGI_10023491), *ARNT* (CGI_10017200), *Cranky* (CGI_10010544), *Mlx2* (CGI_10016301), *SRC* (CGI_10025885), and *SREBP* (CGI_10004108) (Figure 2c&d). Furthermore, genes functioning in “oxidation-reduction process (GO:0055114)” and “chitin metabolic process (GO:0006030)” was also enriched. Among these genes, *tyrosinase-like* (CGI_10007793) possibly involves in the metabolism of tyrosine, at the upstream of epinephrine biosynthesis. Some PIF-like (CGI_10028014, CGI_10004086) and chitin-binding domain containing genes (CGI_10007200, CGI_10012352, CGI_10017942, CGI_10016016) possibly involve in the calcified shell formation.

**Figure 2.**
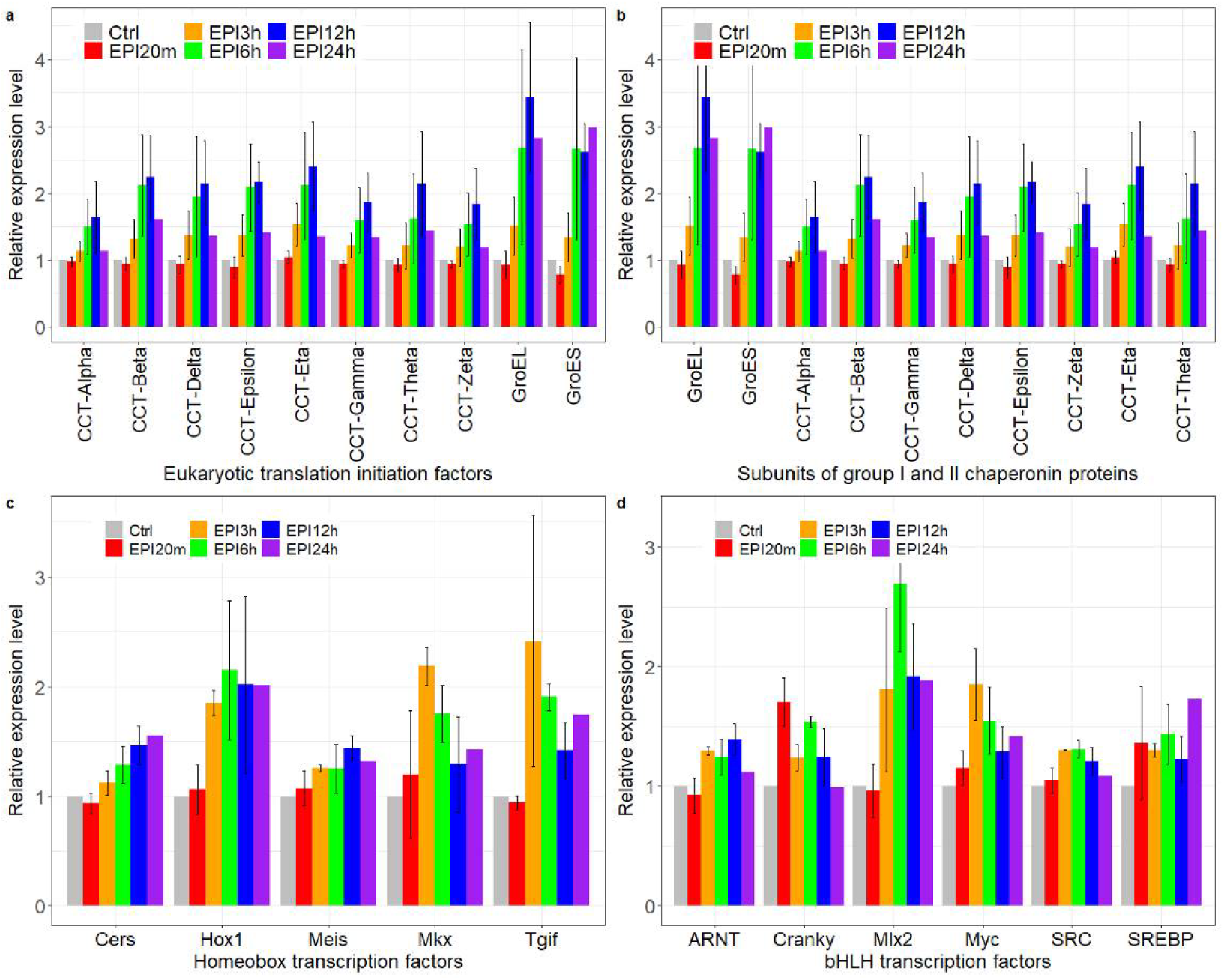
Genes upregulated after EPI stimulation.

GO enrichment analysis on DRGs provided some general terms, e.g., “Protein binding (GO:0005515)”, “microtubule motor activity (GO:0003777)”, “calcium ion binding (GO:0005509)”, “cilium movement (GO:0003341)”, and “calcium-dependent cysteine-type endopeptidase activity (GO:0004198)” (Supplementary Figure S4). Few genes experienced sharp downregulation post epinephrine stimulation. Some pediveliger specifically enriched genes were also down-regulated after epinephrine stimulation: a possible *pacifastin* (CGI_10011175), which inhibits the serine peptidases trypsin and chymotrypsin; a *tyrosinase* (CGI_10006802) and some collagens (CGI_10003774, CGI_10010375, and CGI_10010376). Several transcription factors were also listed in DRGs, such as homeobox gene *Otx* (CGI_10015784), bHLH gene *AP4* (CGI_10026909), forkhead box genes *FoxAB* (CGI_10011631) and *FoxK* (CGI_10026255).

### Intersections between different gene sets

Intersection analysis was conducted on three gene sets: pediveliger enriched genes (389), URGs determined through spat (1390), and EPI (825) experiments. The results indicated a low proportion of overlaps between these gene sets. In the 145 overlapped URGs from both spat and EPI experiments, genes related to “chitin metabolic process” (GO:0006030) were enriched, indicating the initiation of gene regulation for the formation of adult calcified shell. A total of 17 genes were listed in all the three gene sets (Supplementary Figure S5), including three possible shell formation related genes: a *spidroin like glycine-rich protein gene* (CGI_10002595) and two tyrosinase genes (CGI_10017214, CGI_10026227); two genes associated with oxidation-reduction activity: an NADPH oxidase gene (CGI_10025588), a cytochrome P450 gene (CGI_10006946); and a possible D-beta-hydroxybutyrate dehydrogenase gene (CGI_10004113); two stress response genes: a small heat shock protein gene (CGI_10019738), and a gene similar to universal stress protein Usp (CGI_10008265); as well as some other well-annotated genes included an aquaporin-8 (CGI_10018625), a histidine-rich protein (CGI_10016088), a possible serine protease inhibitor (CGI_10010154). As many as 99 (38%) out of the 259 reported shell proteins (Zhang et al., 2012) was listed in either EPI URGs (33 genes) or spat URGs (66 genes), in which 23 were listed in the overlap gene sets (Supplementary Figure S6).

### Proteomic response of pediveliger to epinephrine

Proteome analysis was conducted with twice repeats on two time points post EPI treatment (EPI3h and EPI12h). After the Mascot search, a total of 19,843 unique peptides from 53,056 spectra was identified, corresponding to 5,144 protein groups (Supplementary Figure S7). The length of the identified peptides was around 11 amino acids (Supplementary Figure S8). More than 50% of proteins identified very few peptides, and sequences coverage was lower than 10% (Supplementary Figures S9 & S10). A total of 69 upregulated proteins (URPs) and 33 downregulated proteins (DRPs) were identified for EPI3h samples, 71 URPs, and 55 DRPs for EPI12h samples. Five genes were identified in the overlap of both URPs, including a serine/threonine-protein kinase (CGI_10017094), a pregnancy zone protein (PZP, CGI_10006874), and a 28S ribosomal protein S9 (CGI_10021718). Nine genes showed coincident up-regulation pattern at mRNA expression and protein translation level, including genes related with cellular protein trafficking: a nucleoprotein TPR (CGI_10015574), and a transmembrane emp24 domain-containing protein (CGI_10016770), signal recognition particle 9 kDa protein (SRP, CGI_10010631); genes involving in signal transductions: cAMP-dependent protein kinase type II regulatory subunit (PRKAR2B, CGI_10014742); some enzymes: glutamine synthetase (CGI_10015426), deoxyuridine 5’-triphosphate nucleotidohydrolase (DUT, CGI_10019949); as well as some other genes: sarcalumenin (CGI_10008403), apoptotic chromatin condensation inducer in the nucleus (CGI_10022251), and CDV3-like protein (CGI_10018566).

## Discussion

The life type of oyster transformed from pelagic to benthic through complex metamorphosis. Larvae experience dramatic tissue transition(Coon et al., 1987). Some larval tissues degenerate during metamorphosis, e.g., the ciliated velum, anterior adductor, velum, and ventral retractors (Li et al., 2019). Histogenesis of some adult tissues (e.g., gill (Cannuel et al., 2006) and calcified shells) are initiated during metamorphosis, as well as the formation of pediveliger specific tissues (e.g., foot and pigmented eyespot). Two apoptosis-related genes, *Caspase7* (CGI_10023427) and *Bcl2-like* (CGI_10021873), showed upregulation post epinephrine stimulation, indicating their possible involvement during larval tissues degeneration. However, no significant apoptosis-related genes were identified in pediveliger enriched genes and spat enriched genes. DEGs obtained from epinephrine and pediveliger/spat experiments were distinct inferring from the results of GO enrichment analysis. As marine animals usually conduct rapid metamorphosis (Hadfield, 2000), sampling time point is vital to obtain the metamorphosis related gene set. Pediveliger and spat enriched genes were hard to reflect the gene dynamics during metamorphosis, as the development stages may not overlap with metamorphosis. Epinephrine stimulation on competent pediveligers provides a relatively better way to produce artificially controlled metamorphosis larvae and assay gene expression dynamic during metamorphosis. Together with clues from pediveliger and spat enriched genes, the gene regulation network dynamics during metamorphosis was thus inferred.

### Nervous system remodeling before metamorphosis

As the signal of metamorphosis competent, the appearance of eyespot indicated the possible remodel of the larval nervous system. Foot as a temporary organ to sense a suitable environment for settlement suggests biogenesis of new nerve connections. Correspondingly, genes involved in nervous system development, synapses formation, or neuronal regeneration were highly expressed during the pediveliger stage: *SCO-spondins* (CGI_10022619 & CGI_10022620), *neuroligins* (CGI_10007153, CGI_10007154), *neurotrimin* (CGI_10014425) and *tenascin* (CGI_10002258). At the same time, expression levels of many receptors for sensing environmental stimuli were also upregulated, together with some neuropeptide receptor genes. Epinephrine, gamma-aminobutyric acid (GABA), dopamine, and acetylcholine have been proved to be effective neuroendocrine compounds to induce oyster metamorphosis (Joyce et al., 2018). There possible receptors all upregulated during the pediveliger stage (see “specifically enriched genes during the pediveliger stage” in the Result part). However, little is known on their functional pathway. Take an *octopamine receptor* gene (OAR, CGI_10017568), for example, which was also upregulated post epinephrine stimulation. This gene was homology to *Lymnaea* OAR2 and should belong to a poorly studied subfamily of invertebrate OA/TA receptors. The protein distributes at the surface of the pediveliger foot, indicating the possible involvement to sense environmental stimuli. However, direct treatment of competent pediveliger with octopamine resulted in a weak effect to induce metamorphosis. Octopamine also showed no effect on the cAMP or Ca^2+^ signal pathway when treating HEK293 cells stably expressing this gene (Ji et al., 2016). Functional assays to these key receptors will provide further insight into their molecular evolution, as well as help to develop new strategies to control the settlement of marine animal metamorphosis more effectively.

### Shell formation

Oyster shell experience transformation during metamorphosis (Figure 1a), from aragonite larval shell to calcite adult shell (Stenzel, 1964, Haley et al., 2018). This was well reflected by gene expression dynamics revealed through Venn analysis on three gene sets: pediveliger enriched genes (389), URGs determined through spat (1390), and EPI (825) experiments. In the 145 overlapped URGs from both spat and EPI experiments, only the term “chitin metabolic process” (GO:0006030) was enriched, indicating the acute response of genes related to shell formation. For example, gigasin-6, which was identified from oyster calcifying matrix (Marie et al., 2011), expressed higher after EPI stimuli and post attachment. It almost specifically express in the mantle and possibly function during shell calcifying. As many as 99 (38%) out of the 259 reported adult shell proteins (Zhang et al., 2012) was listed in either EPI URGs (33 genes) or spat URGs (89 genes), in which 23 were listed in the overlap gene sets. As adult oyster shells still have small areas of aragonite (Stenzel, 1963), which usually stacks with chitin sheets and Pif complexes (Pif 97, Pif 80, N16, and other proteins) to form nacreous layer(Suzuki et al., 2009). Accordingly, some *PIF-like* genes and *chitin-binding domain containing protein* genes were upregulated post EPI stimulation (see the result part “Transcriptomic response of pediveligers to epinephrine”).

### Global activation of protein translation initiation system

Protein synthesis is principally regulated at the initiation stage (Jackson et al., 2010). EPI stimulation on oyster pediveliger upregulated the expression level of five (eIF1, eIF1A, eIF2, eIF3, and eIF5) out of the nine core eukaryotic translation initiation factors (plus eIF4A, eIF4B, eIF4F, and eIF5B) (Jackson et al., 2010). Correspondingly, the expression level of genes coding chaperones and chaperonin proteins which facilitate protein folding or activated proteolysis of misfolded proteins (Slavotinek et al., 2001) were also upregulated (Figure 3). Within the nine genes, which have been detected to be up-regulated from both transcriptome and proteome results, three are related with cytoplasmic trafficking of mRNA or protein: nucleoprotein *TPR* gene (CGI_10015574) which involves in the trafficking of proteins and mRNA across nuclear and cytoplasm; *transmembrane emp24 domain-containing protein* (CGI_10016770) which involves in intracellular protein transport; *signal recognition particle 9 kDa protein* (CGI_10010631) which plays an important role in targeting secretory proteins to the rough endoplasmic reticulum membrane.

**Figure 3.**
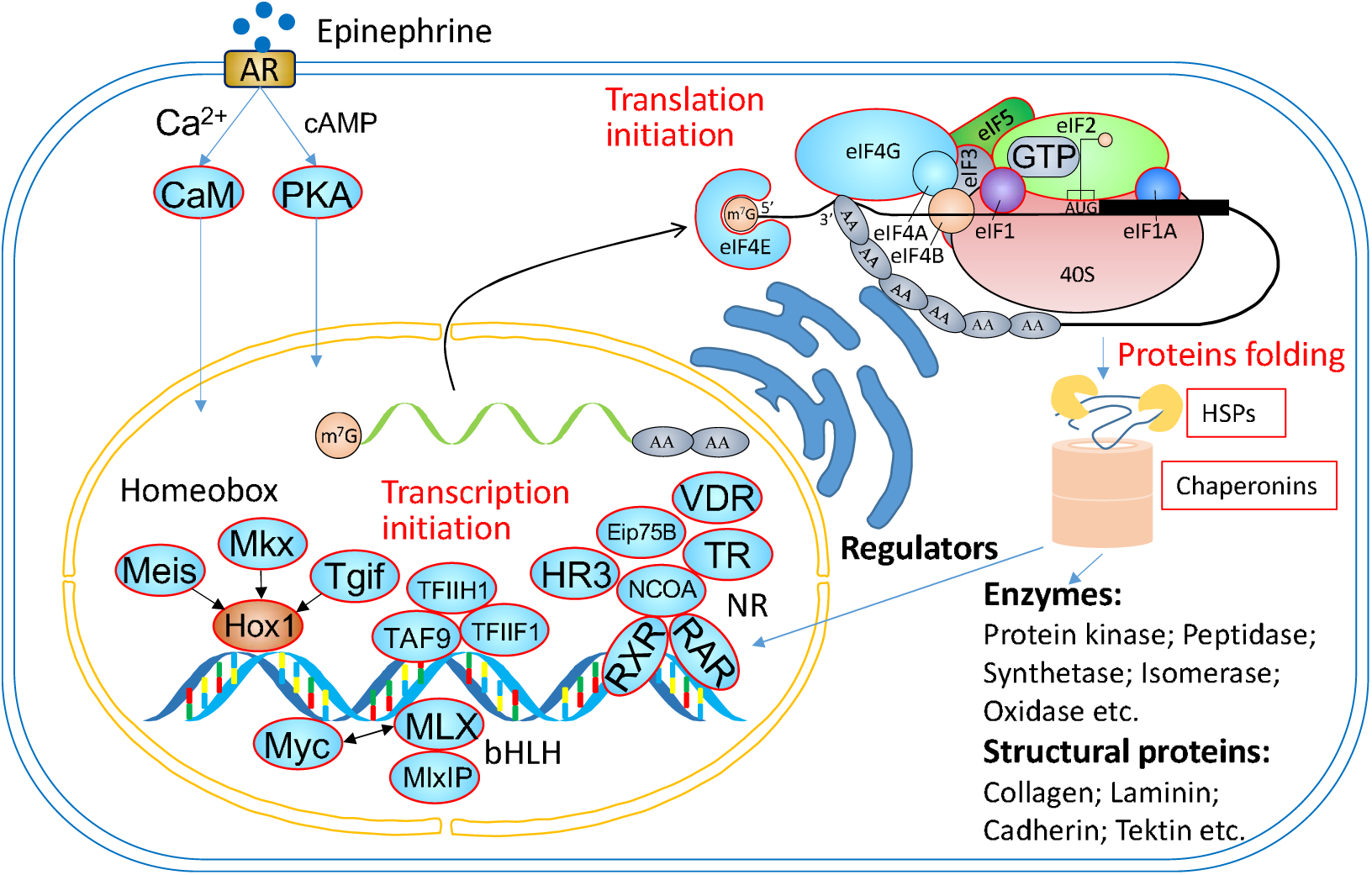
Gene response of oyster competent pediveliger to epinephrine. Shapes with red edge denote upregulated genes (URGs). Abbreviations: AR, adrenergic receptor; CaM, calmodulin; PKA, cAMP-dependent protein kinase; Meis, homeobox protein Meis; Mkx, homeobox protein Mohawk; Tgif, TGFB induced factor homeobox; Hox1, homeobox 1; TFIIH1, general transcription factor IIH, polypeptide 1, 62kDa-like; TFIIF1, General transcription factor IIF subunit 1; TAF9, TATA-Box binding protein associated factor 9; bHLH, basic helix-loop-helix transcription factors; Myc, MYC proto-oncogene; MLX, MAX dimerization protein MLX; MlxIP, MLX interacting protein; NR, nuclear receptor; VDR, vitamin D receptor; THR, thyroid hormone receptor; NCOA, nuclear receptor coactivator; RXR, retinoid X receptor; RAR, retinoic acid receptor; Eip75B, Ecdysone-induced protein 75B; HR3, nuclear hormone receptor HR3; eIF, Eukaryotic translation initiation factor; HSP, heat shock protein.

Furthermore, expression levels of many transcription factors were also upregulated. Expression increasing of the nuclear receptors (NRs), as well as their interacting partners RXR and RAR, indicated the possible involvement of nonpolar regulatory molecules in downstream gene transcription post epinephrine stimulation. These NRs included potential homologs that play essential roles during insect metamorphosis, e.g., Ecdysone-induced protein 75B (Guo et al., 2016) (Eip75B, CGI_10004075 in the oyster), nuclear hormone receptor HR3 (Zhao et al., 2018) (CGI_10011713 in the oyster). Both genes respond to the steroid hormone 20-Hydroxyecdysone (20E) and its signaling pathway in *D. melanogaster*. The widespread of ecdysteroids have been reported in many molluscs (Romer, 1979, Whitehead et al., 1982, Mukai et al., 2001) and other Lophtrochozoans (Lafont et al., 2009). However, the physiological function remains to be studied. At the same time, another upregulated NR, vitamin D receptor (VDR, CGI_10020896 in the oyster), indicated the possible regulation dynamic of calcium homeostasis control during metamorphosis. Different temporal patterns of response were also observed in the DEGs, indicating the hierarchical model of gene regulation. Take the EPI responsive Homeobox and bHLH genes, for example, bHLH genes *Cranky, Myc*, and *SREBP* had a quick response and increased expression level within 20 minutes post EPI stimulation, while *Hox1, Mkx*, and *Tgif* homeobox genes and *ARNT, Mlx2, SRC* bHLH genes response within 3 hours. *Cers* and *Meis* homeobox genes upregulated gradually within 24 hours. Although 20 minutes of EPI treatment was enough to initiate the metamorphosis and should have triggered the genic response in the competent oyster larvae, the genomic-scale gene expression response did not occur within as short as 20 minutes. PCA analysis indicated that the comprehensive gene expression response should have occurred between 20 minutes to 3 hours.

There were also transcription factors downregulated during metamorphosis, among which, turning off of gene FoxAB (CGI_10011631) can be a biomarker for the start of oyster metamorphosis transforming. FoxAB was firstly identified in the sea urchin genome (Tu et al., 2006). It is an ancient family but has been lost several times during the Bilateria evolution. None member from this family has been identified from vertebrates and ecdysozoans (Tu et al., 2006, Yu et al., 2008, Yang et al., 2014). Species whose FoxAB gene have been reported include the sea urchin *Strongylocentrotus purpuratus* (Tu et al., 2006), the cephalochordate amphioxus *Branchiostoma floridae* (Yu et al., 2008), the enteropneust hemichordate *Saccoglossus kowalevskii* (Fritzenwanker et al., 2014), hydrozoan cnidarian *Clytia hemisphaerica* (Chevalier et al., 2006), the oyster *C. gigas*, the limpet *Lottia gigantean*, the annelid *Capitella teleta* (Boyle et al., 2014, Yang et al., 2014), and the bryozoan *Bugula neritina* (Fuchs et al., 2011). The developmental expression pattern of *FoxAB* has been well studied in the last species, where *BnFoxAB* was also exclusively expressed in larval tissues and discarded during metamorphosis (Fuchs et al., 2011). Knowledge of the exact function of FoxAB is still limited, but it is possibly recruited to involve in the formation of ectoderm originated larval tissues, such as oral ectoderm (Boyle et al., 2014).

## Conclusion

Gene response of the Pacific oyster competent pediveliger to epinephrine was assayed by RNAseq and proteomic analysis. By integrating the pediveliger and spat enriched gene sets, gene network dynamics during oyster metamorphosis was dissected. Different gene regulation patterns were observed for epinephrine responsive gene set, pediveliger, and spat enriched gene sets, indicating quick changing of gene regulation dynamics. During the pediveliger stage, genes related to the integral component of membrane (receptors) and nervous system formation were massively upregulated, indicating structural preparation for the initiation of metamorphosis. Metamorphosis was quickly simulated post epinephrine treatment on pediveligers, almost the whole component related to protein translation initiation and protein folding control system were upregulated, indicating massive biogenesis. Different transcription factors, including nuclear receptors in ecdysteroids signaling pathways, showed various response patterns to epinephrine stimulation. Structural proteins related to shell formation was upregulated during metamorphosis. Calcified adult shell grew quickly within 24 hours post epinephrine stimulation, and the spat shell size was even doubled based on the larval shell size. Early spat post attachment enriched genes mainly related to adaptation to the benthic environment, such as genes with antioxidation activities. Lophotrochozans represented a large clade of Bilateria and is the key to understand animal evolution. However, there is limited knowledge of the developmental function of organic molecules and genes in Lophotrochozans, such as ecdysteroids and FoxAB. More studies are needed to understand the gene regulation network of Lophotrochozans further.

## Data availability

RNAseq data has been deposited with GenBank under BioProject PRJNA553079. The datasets used and analyzed during the current study are also available from the corresponding author on reasonable request.

## Acknowledgement

We acknowledge the Qingdao Frontier Ocean Seed Company Ltd Company for providing the facility for culturing the oysters. We thank Wen Huang for assistance during the oyster treatment experiment. Most of the calculations were performed using the supercomputer cluster of the High Performance Computing Center (HPCC) at the Institute of Oceanology, Chinese Academy of Sciences. We acknowledge the grant support from the Pilot National Laboratory for Marine Science and Technology (Qingdao) (Grant No. QNLM2016ORP0408), the Key Deployment Project of Centre for Ocean Mega-Research of Science, Chinese Academy of Science (COMS2019Q11), and the National Natural Science Foundation of China (41776152, 31402285).

## Supplementary Figures

**Figure S1.**
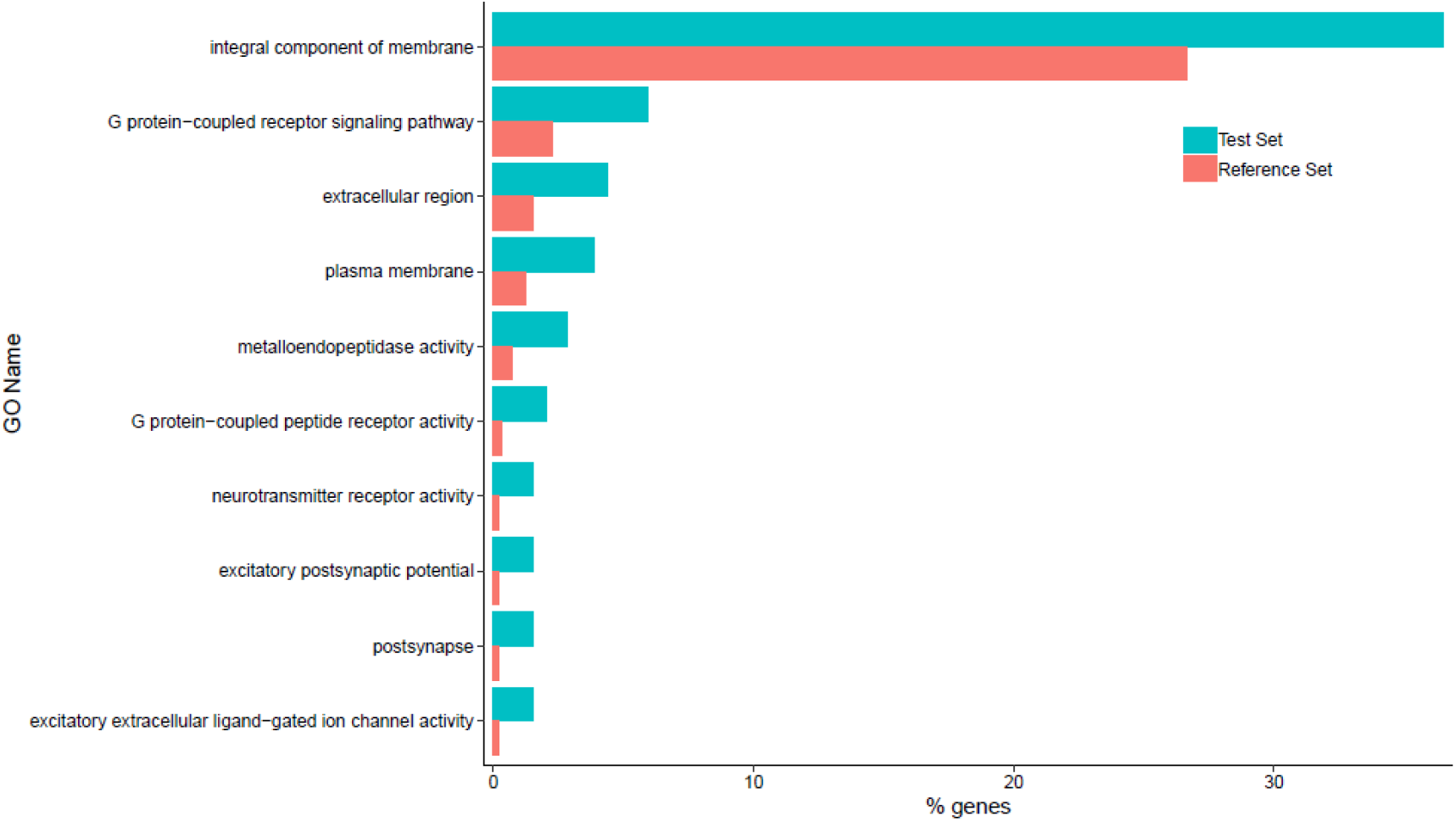
Gene ontology enrichment analysis result of specifically enriched genes during pediveliger stage.

**Figure S2.**
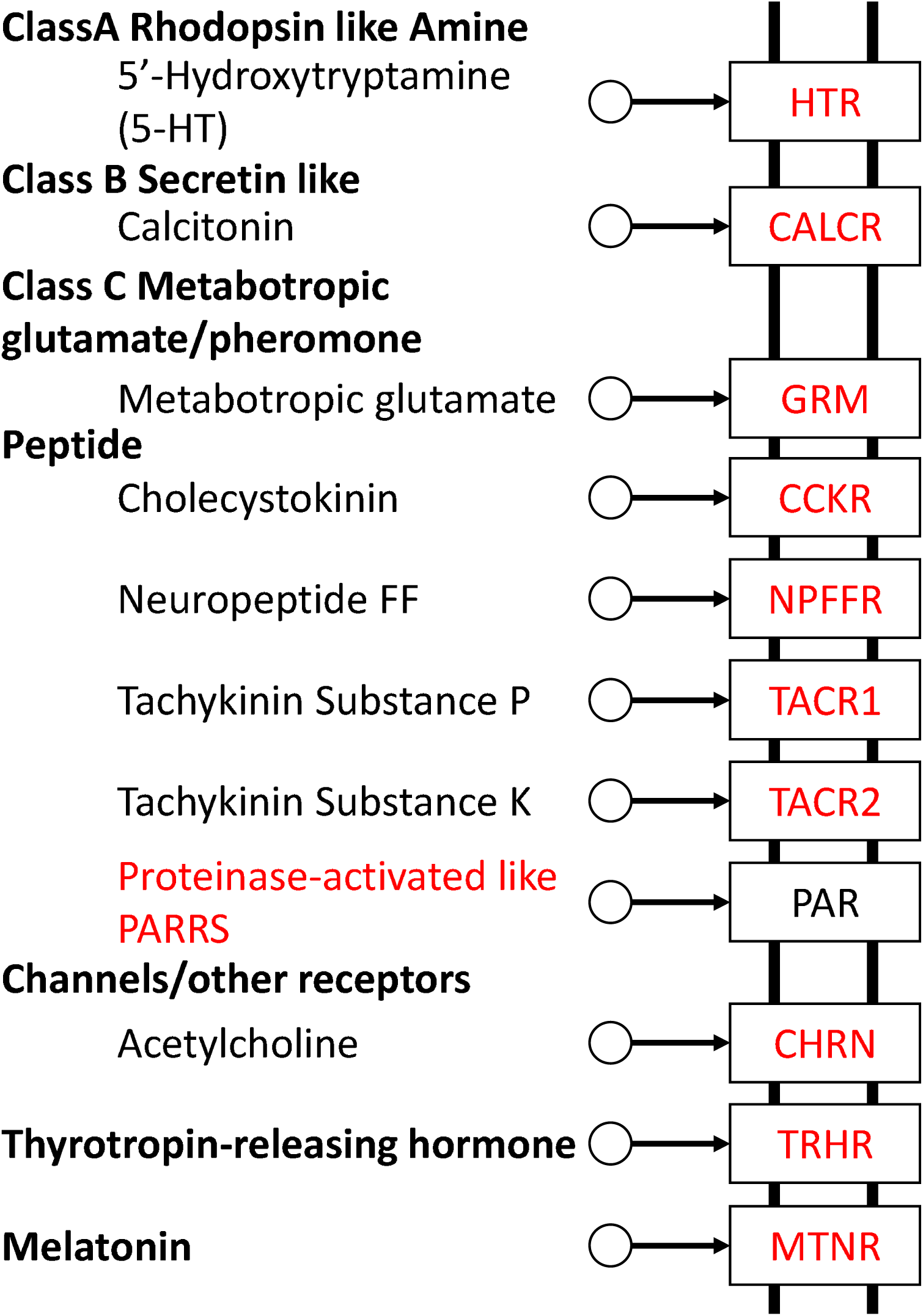
Early spat upregulated genes (red) in the neuroactive ligand-receptor interaction pathway. HTR, 5-hydroxytryptamine receptor; CALCR, calcitonin receptor; GRM, metabotropic glutamate receptor; CCKR, cholecystokinin A receptor; NPFFR, neuropeptide FF receptor; TACR1, tachykinin receptor 1; TACR2, tachykinin receptor 2; CHRN, nicotinic acetylcholine receptor; TRHR, thyrotropin-releasing hormone receptor; MTNR, melatonin receptor.

**Figure S3.**
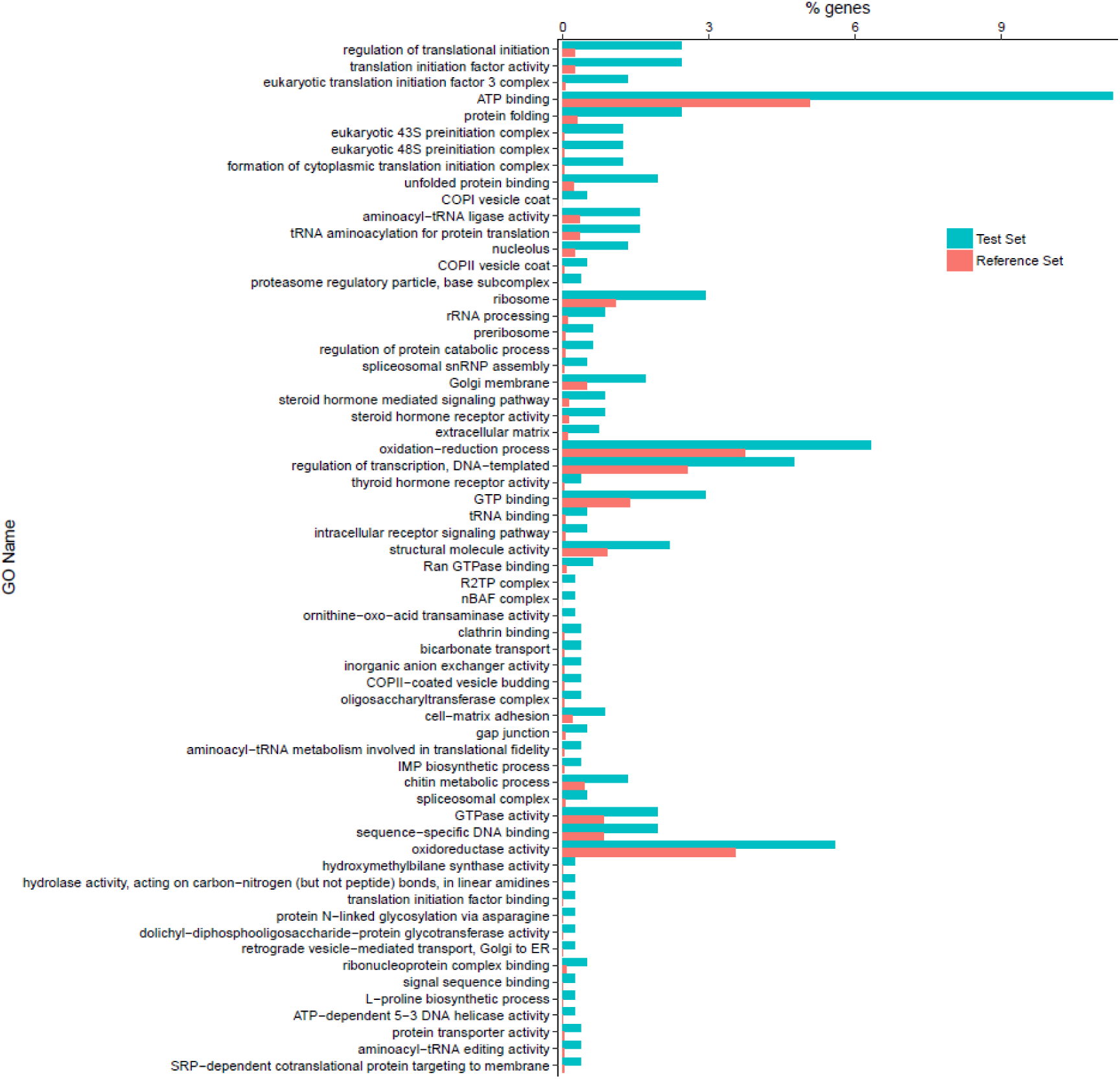
Gene ontology enrichment analysis result of upregulated genes post epinephrine treatment on oyster competent pediveliger.

**Figure S4.**
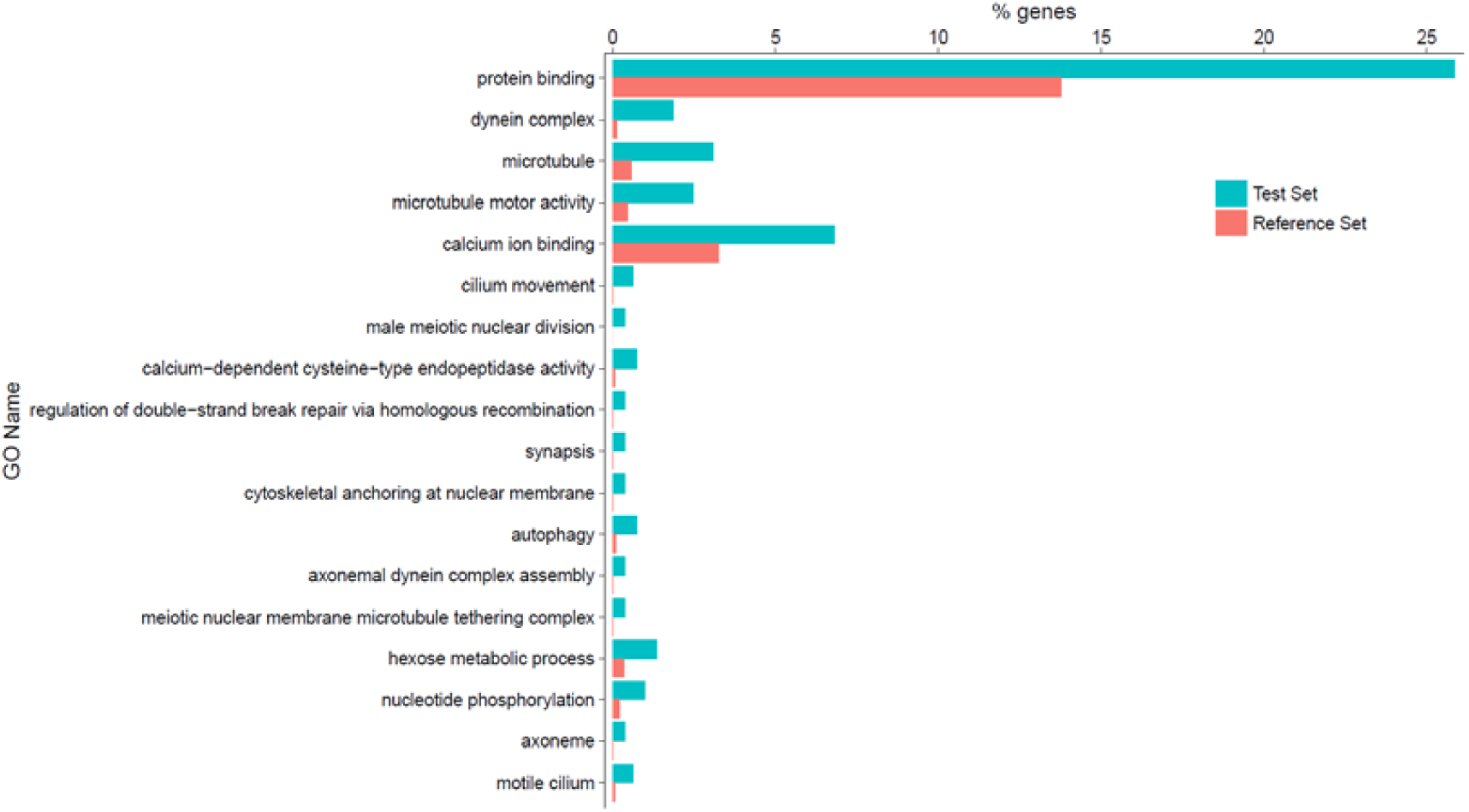
Gene ontology enrichment analysis result of downregulated genes post epinephrine treatment on oyster competent pediveliger.

**Figure S5.**
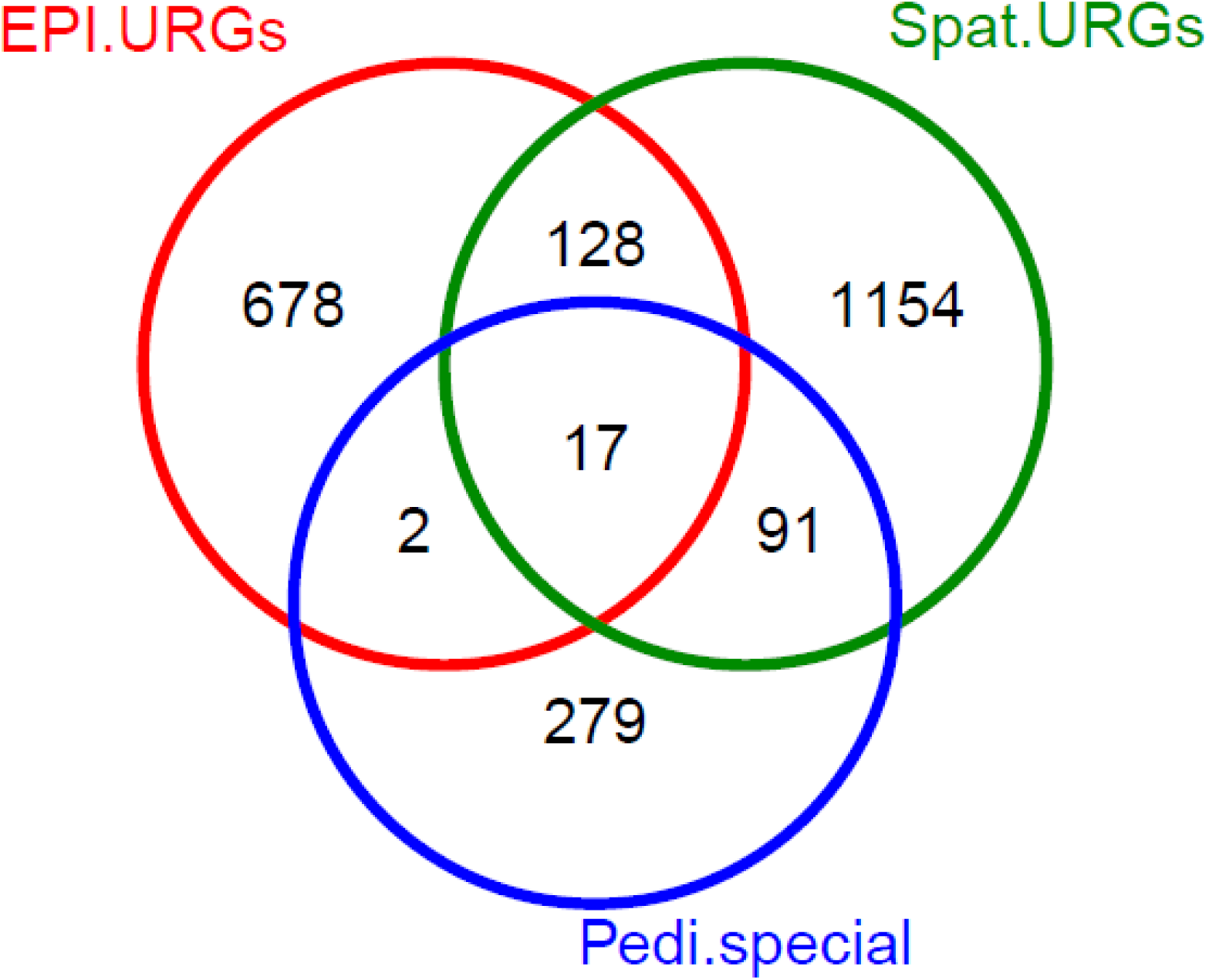
Intersection analysis on three gene sets: pediveliger enriched genes (389), URGs determined through spat (1390) and EPI (825) experiments.

**Figure S6.**
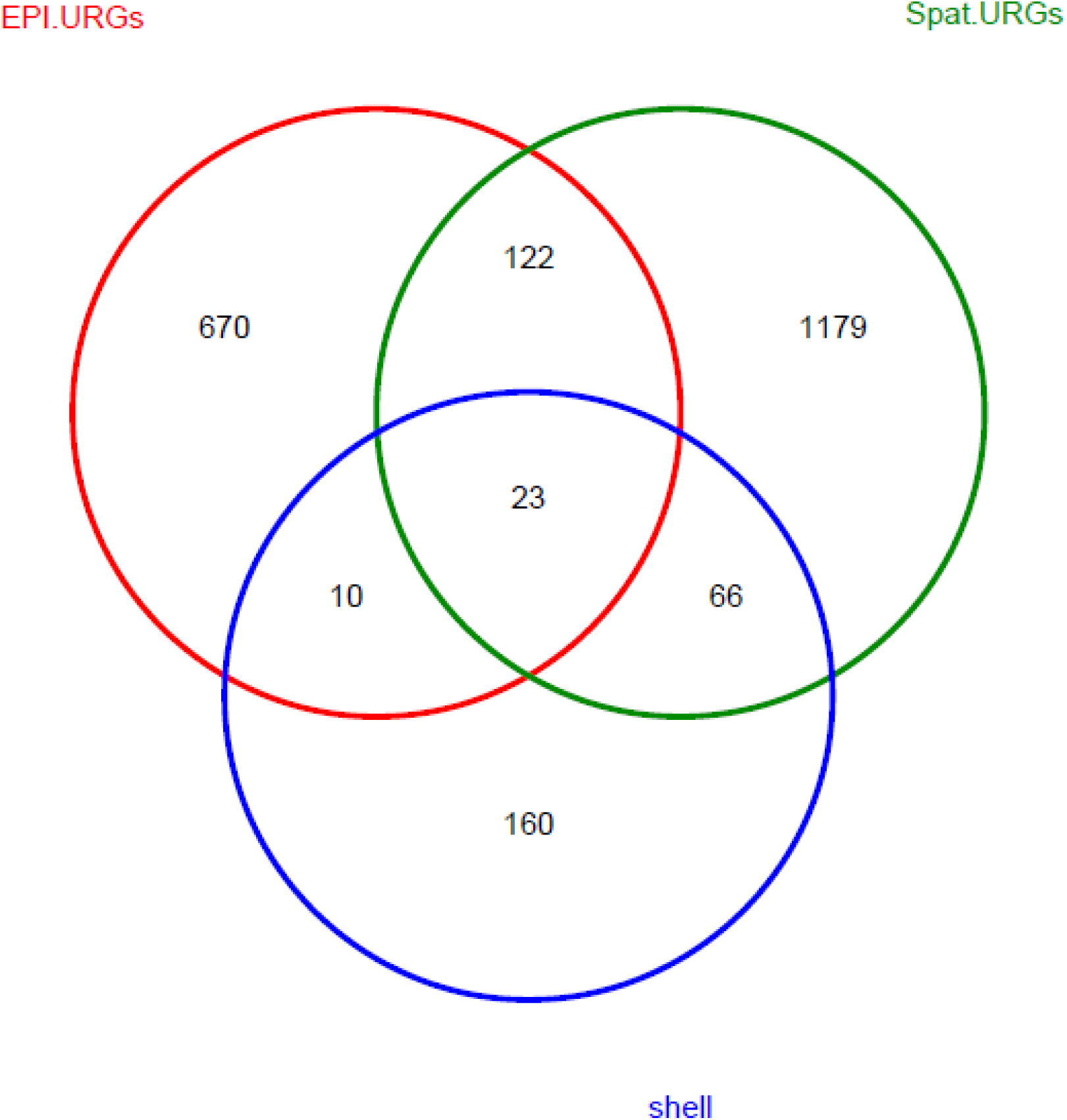
Intersection analysis on three gene sets: shell identified protein genes (259), URGs determined through spat (1390) and EPI (825) experiments.

**Figure S7.**
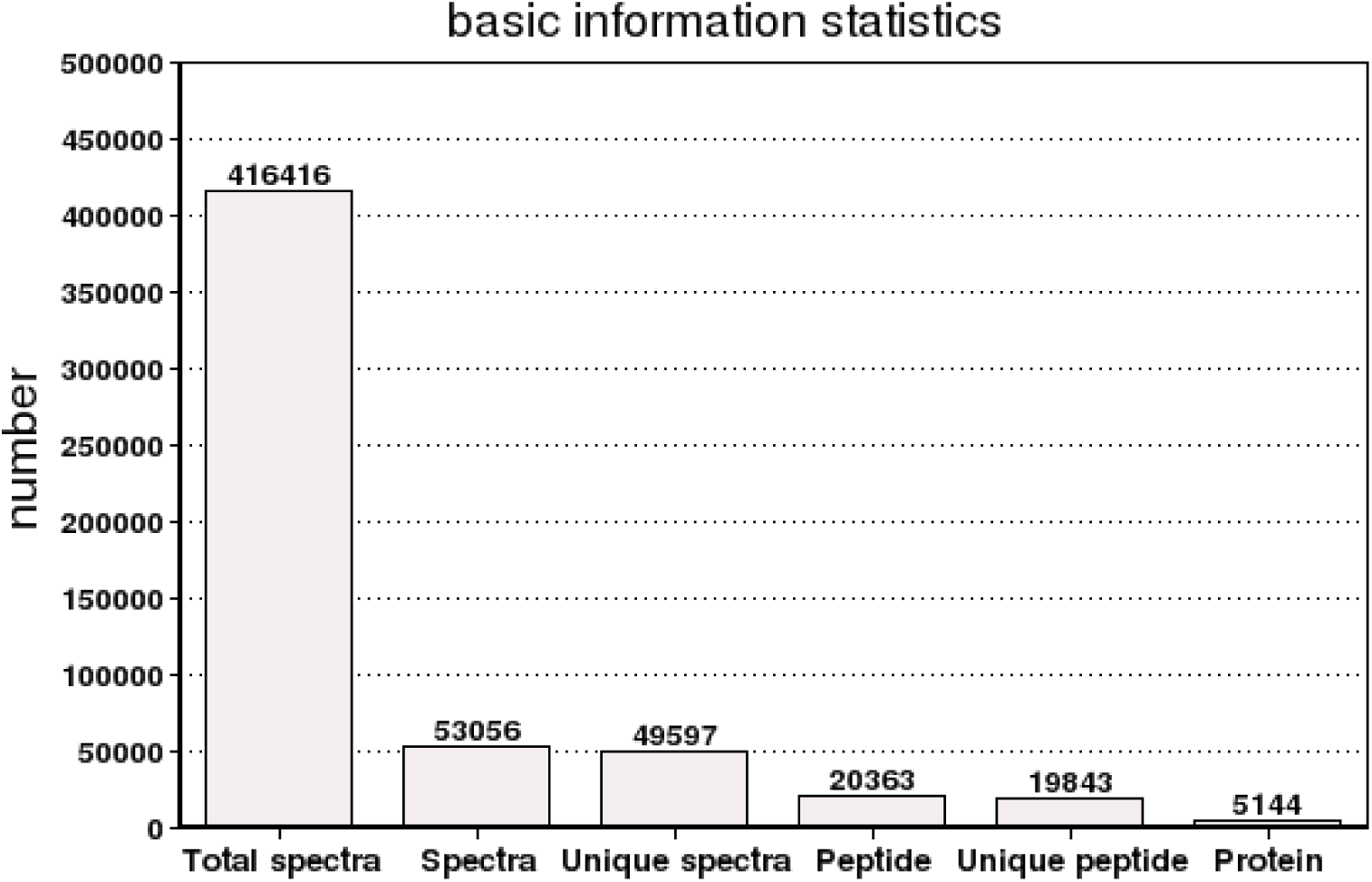
Summary of proteome identification with iTRAQ analysis. Total spectra is the total number counted by secondary spectrum. Spectra indicate the number of successfully matched. Unique spectra indicate the number successfully matched unique peptides. Peptide is the number of identified peptide. Unique peptide indicates the number of identified unique peptides. Protein indicates the total number of identified proteins.

**Figure S8.**
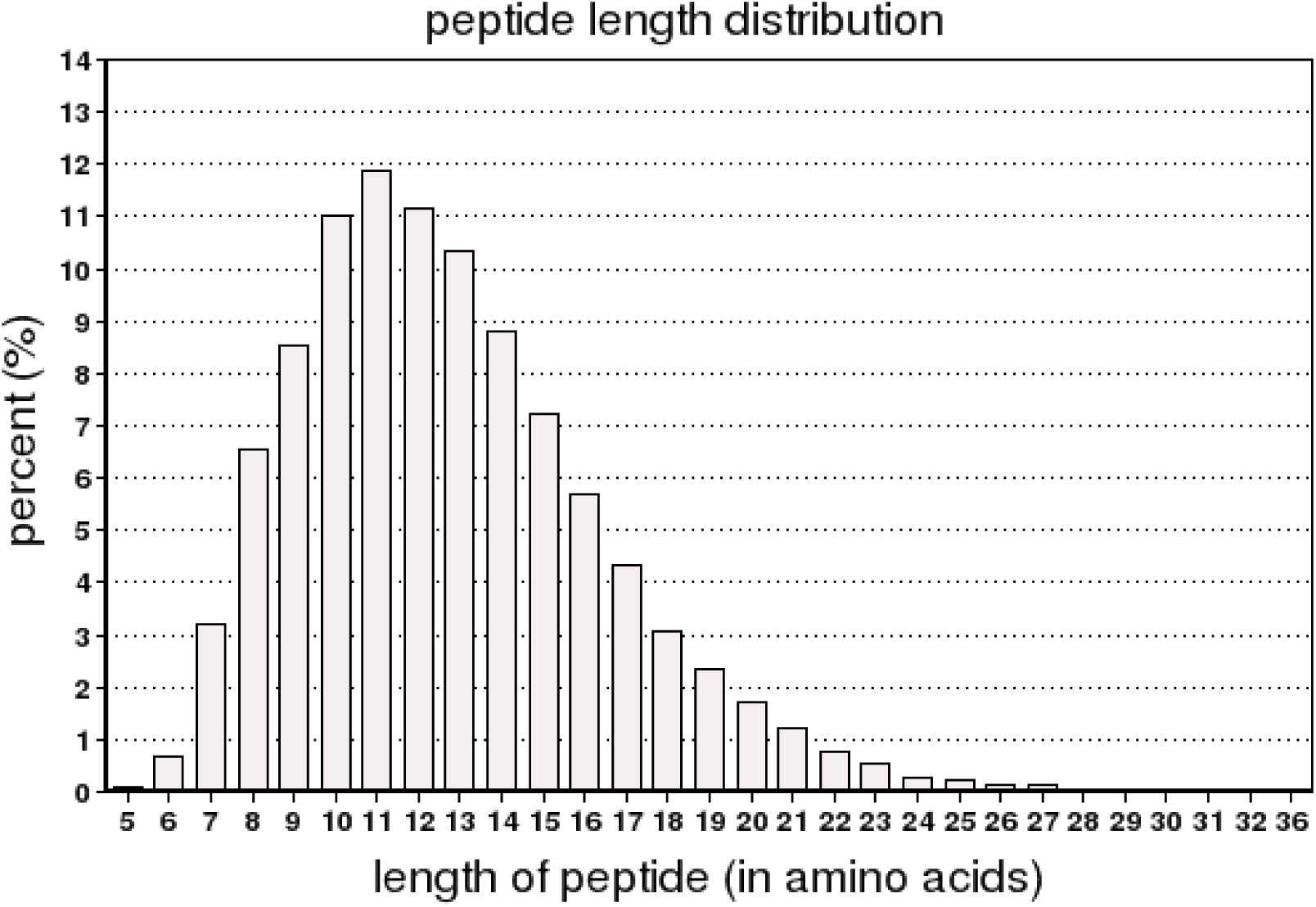
Distribution of the length of identified peptides.

**Figure S9.**
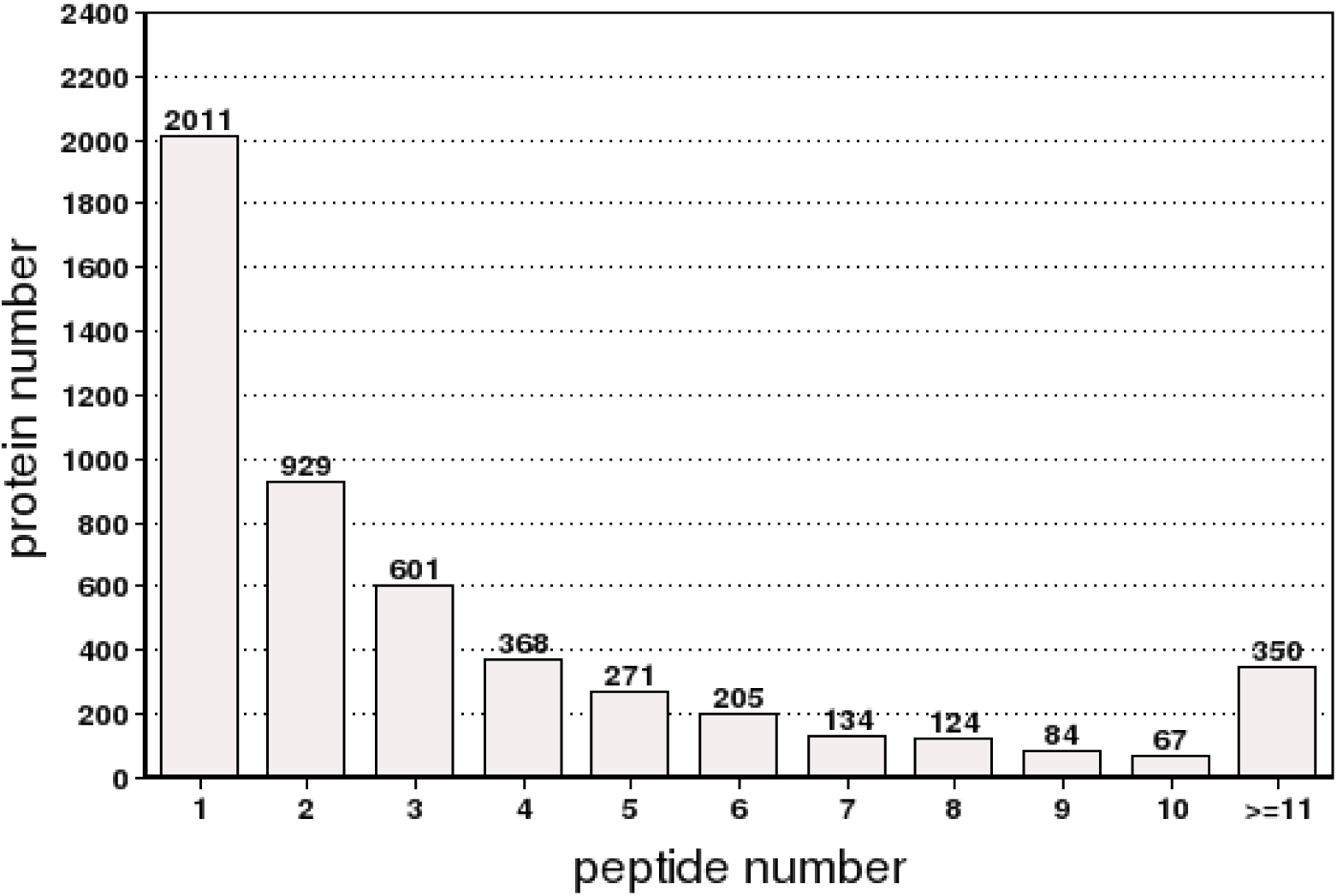
Distribution of peptide number.

**Figure S10.**
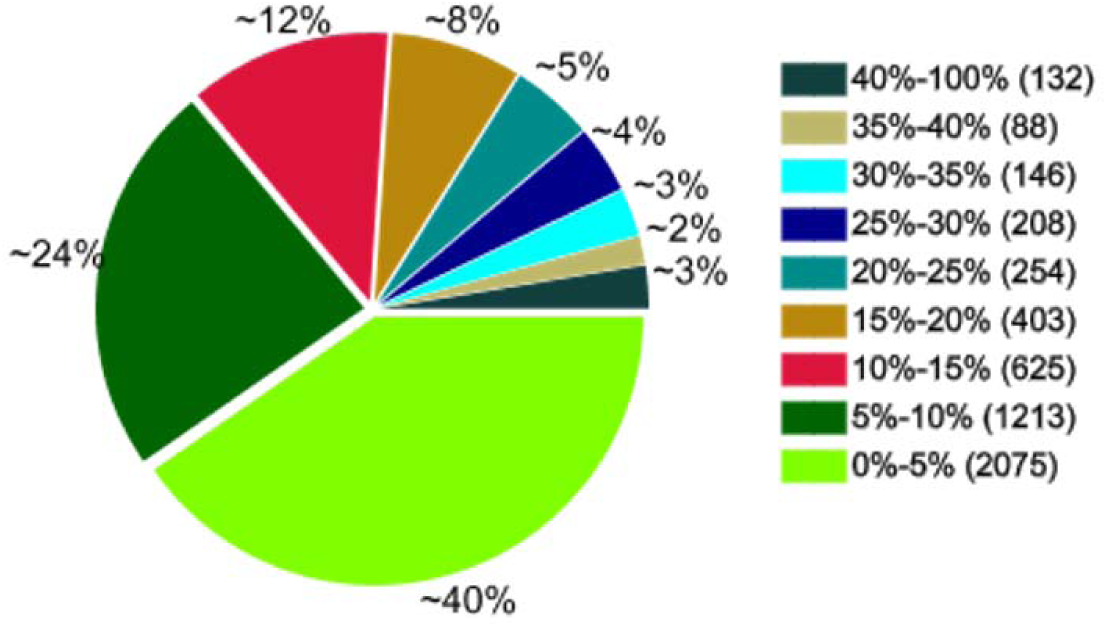
Distribution of protein’s sequences coverage by their identified peptides.

